# IL1A enhances TNF-induced retinal ganglion cell death

**DOI:** 10.1101/2024.05.28.596328

**Authors:** Katherine M. Andersh, Michael MacLean, Gareth R. Howell, Richard T. Libby

**Affiliations:** Department of Ophthalmology, Flaum Eye Institute, University of Rochester Medical Center, Rochester, NY, USA; Neuroscience Graduate Program, University of Rochester Medical Center, Rochester, NY, USA; The Center for Visual Sciences, University of Rochester, Rochester, NY, USA; Department of Biomedical Genetics, University of Rochester Medical Center, Rochester, NY, USA; The Jackson Laboratory, 600 Main Street, Bar Harbor, ME, USA

**Keywords:** proinflammatory cytokines, interleukin 1, neuroinflammation, glaucoma, optic neuropathy

## Abstract

Glaucoma is a neurodegenerative disease that leads to the death of retinal ganglion cells (RGCs). A growing body of literature suggests a role for neuroinflammation in RGC death after glaucoma-relevant insults. For instance, it was shown that deficiency of three proinflammatory cytokines, complement component 1, subcomponent q (*C1q*), interleukin 1 alpha (*Il1a*), and tumor necrosis factor (*Tnf*), resulted in near complete protection of RGCs after two glaucoma-relevant insults, optic nerve injury and ocular hypertension. While TNF and C1Q have been extensively investigated in glaucoma-relevant model systems, the role of IL1A in RGC is not as well defined. Thus, we investigated the direct neurotoxicity of IL1A on RGCs in vivo. Intravitreal injection of IL1A did not result in RGC death at either 14 days or 12 weeks after insult. Consistent with previous studies, TNF injection did not result in significant RGC loss at 14 days but did after 12 weeks. Interestingly, IL1A+TNF resulted in a relatively rapid RGC death, driving significant RGC loss two weeks after injection. JUN activation and SARM1 have been implicated in RGC death in glaucoma and after cytokine insult. Using mice deficient in JUN or SARM1, we show RGC loss after IL1A+TNF insult is JUN-independent and SARM1-dependent. Furthermore, RNA-seq analysis showed that RGC death by SARM1 deficiency does not stop the neuroinflammatory response to IL1A+TNF. These findings indicate that IL1A can potentiate TNF-induced RGC death after combined insult is likely driven by a SARM1-dependent RGC intrinsic signaling pathway.

## Introduction

Glaucoma is a neurodegenerative disease characterized by retinal ganglion cell (RGC) death and irreversible loss of vision. Many studies in recent years have identified RGC intrinsic pathways required for RGC death after glaucoma-relevant insults ^1^. While these findings are promising, recent studies have implicated extrinsic signaling as key drivers of RGC injury in glaucoma. In fact, multiple studies have suggested a primary or direct role of microglia and astrocytes in glaucomatous neurodegeneration ^2, 3^.

Recent studies showed a direct neurotoxic role of neuroinflammation in glaucoma relevant RGC death, specifically through 3 neuroinflammatory molecules: *C1qa*, *Tnf*, and *Il1a* ^4^. These molecules were capable of indirectly driving RGC death via astrocyte activation *in vitro* ^4^, and deletion of *C1qa*, *Tnf*, and *Il1a in vivo* was sufficient to protect RGCs from mechanical optic nerve injury (optic nerve crush; ONC) ^4, 5^ and ocular hypertension (OHT) ^6^. These studies show the role of neuroinflammatory signaling in driving glaucoma-relevant RGC death. In addition, *in vitro* application of TNF, IL1A and C1Q, can drive astrocyte activation ^4^. IL1A, TNF, and C1Q exposed astrocytes release neurotoxic lipids, which can kill RGCs ^7^. While these findings demonstrate the importance of these molecules in RGC degeneration after axonal insult, the direct neurotoxicity of these molecules in the absence of other insult(s) remains understudied.

C1Q and TNF have been extensively studied in the context of glaucoma, with *C1q* inhibition or deletion proving to be partially protective to RGCs^8^. However, conflicting results implicating TNF in RGC degeneration have made it unclear whether it is protective or detrimental. TNF inhibition results in mild protection from OHT ^9, 10^, and deletion of *Tnf* leads to worsening RGC degeneration after ONC ^11^. In the absence of other injuries, direct TNF application is known to drive approximately 15-20% RGC loss at a delayed rate following 8-12 weeks ^11, 12, 13^. These data suggest a complex and context-dependent role of TNF. IL1A is another key proinflammatory cytokine capable of being released from several cell types, including microglia ^14, 15, 16^, astrocytes ^17, 18, 19, 20^, and Müller glia ^21^. While IL1A was shown to be expressed in early stages of glaucoma in the DBA/2J glaucoma model ^8^, study of its function in the retina has been limited to retinal ischemia^22^ and uveitis ^23, 24^. Several studies assess the neurotoxic effects of IL1A in the central nervous system ^20, 25, 26, 27^ though none have focused on the retina. It remains unknown if and how IL1A may act to drive RGC degeneration. Given the role of IL1A in RGC death after glaucoma-relevant insults learned from loss of function studies ^4, 5, 6, 28^ it is important to understand the effects of IL1A on RGC survival. Here, we directly test the role of IL1A on RGCs to gain insight into the mechanisms of RGC death after cytokine insult in the absence of other injuries.

## Methods

### Mice

Animals were fed chow and water ad libitum and housed on a 12-hour light-to dark cycle. Roughly equal numbers of males and females were used for each experimental group. All mice included were 2–6 months of age. C57BL/6J, *Sarm1* deficient (Jackson Laboratory, Stock # 018069), or *Jun* floxed (*Jun^fl^*) ^29^ mice were used. Floxed alleles of *Jun* were recombined in RGCs Six3-cre (The Jackson Laboratory, Stock# 019755) ^30^. All experiments were conducted in adherence with the Association for Research in Vision and Ophthalmology’s statement on the use of animals in ophthalmic and vision research and were approved by the University of Rochester’s University Committee on Animal Resources.

### Intravitreal Injections

Mice were anesthetized with an intraperitoneal injection of 0.05 ml/10 g solution containing ketamine (20 mg/mL) and xylazine (2 mg/ mL). Eyes were sterilized with 50% betadine solution in phosphate buffered saline (PBS). The conjunctiva was cleared away with the bevel of a 30-gauge needle on the temporal side, and a small poke was made with the needle below the limbus and through the sclera. Vitreous was allowed to drain and was wicked away using a Kim wipe. If the eyes experienced an excessive bleed following the initial insertion of the needle, the eye was excluded from further injection and investigation. Hamilton syringes (Hamilton Company, 7633– 01) with blunt 33-gauge needles were used to perform intravitreal injections. The needle of the Hamilton syringe was inserted approximately 1mm into the incision site at a 45° angle toward the optic nerve over a period of 30 seconds. Care was taken to avoid contacting the lens with the Hamilton needle. Before experiments were performed, it was established that eyes with observable lens damage due of intravitreal injection would be excluded from the study. Compounds were diluted in sterile PBS and injected slowly into the vitreous over 2 minutes to prevent a sudden pressure increase or leakage. 2 μL of IL1A alone (0.1 μg/μL, Peprotech, 200-01A), TNF alone (0.05 μg/μL, Sigma, T7539), or IL1A+TNF were injected. After injection, the needle was held in place for 60 seconds to reduce drainage, and the needle was removed from the eye over the course of 30 seconds. The contralateral eye was injected with 2 μL of sterile PBS with identical methods to serve as an internal control. These methods were consistent across all lines and genotypes used.

### Immunostaining and Quantification

Immunohistochemistry was performed as previously described ^31, 32, 33, 34, 35^. Eyes were enucleated and fixed in 4% paraformaldehyde in 1X PBS at room temperature for 2 hours. For whole mount assessment, retinas were dissected from the optic cup and blocked in 10% horse serum, 0.4% TritonX in 1X PBS overnight at 4 °C. Retinas were incubated at 4 °C for 72 h in primary antibody diluted in 10% horse serum, 0.4% TritonX in 1X PBS. Primary antibodies included: rabbit anti-pJUN (phosorylated JUN, Cell Signaling, 1:250; Cat# 9261S), rabbit cCASP3 (cleaved caspase 3, R&D, 1:500), rabbit anti-RBPMS (RNA binding protein, mRNA processing factor, GeneTex, 1:500; Cat# GTX118619), guinea pig anti-RBPMS (Phosphosolutions, 1:500; Cat# 1832-RBPMS), mouse anti-TUJ1 (class III beta-tubulin, BioLegend, 1:1000; cat# 801201). Retinas were then washed and incubated overnight at 4 °C in secondary antibody diluted in PBS (Alexa-fluor conjugated, Invitrogen and JacksonImmuno). Retinas were washed and mounted ganglion cell layer up in Fluorogel in TRIS buffer (Electron Microscopy Sciences). As previously described RBPMS+ and pJUN+ cells were quantified following collection of eight 40x fields per retina, and cCASP3+and RBPMS+ double positive cells were quantified using eight 20x fields per retina. Images were equally spaced 220 μm from the peripheral edge of the retina (with 2 images per quadrant) to gauge gross changes in RGC density following injection. For assessment of RGC density changes following injections, RBPMS+ TUJ1+ RGCs were quantified following collection of eight 40x fields per retina. Image quantification was performed using the cell-counter tool in ImageJ. A masked observer captured and manually quantified all images using these methods across groups and conditions.

### Statistical Analysis

The animal number is provided in each figure legend and, where appropriate, was determined by power analysis. Male and female mice were used for all experiments. Except the RNA-seq data, bar graphs represent means and error bars are standard error of the mean (SEM). For all experiments, mice were randomly chosen for experimental groups and the experimenter was masked to genotype and/or condition. Also, prior to injection, any animals with pre-existing abnormal eye phenotypes (e.g. displaced pupil, cataracts, abnormal size, missing eyes, etc.) were excluded from the study. For calculation of RBPMS+ and pJUN+ cells, pJUN+ cells and total RBPMS+ cells per image were assessed to determine both density of pJUN+ cells and percentage of RBPMS+ pJUN+ double positive RGCs. These data were then assessed for differences using a t-test. Quantification of cCASP3+ RBPMS+ double positive cells was done across all timepoints, separating out IL1A+TNF injected and PBS injected eyes per day, and assessed using a two-way ANOVA. For quantification of RGC density, RBPMS+ and TUJ1+ double positive cells were counted in all images per group and compared using a t-test for those passing the Shapiro Wilk normality test (IL1A+TNF versus PBS), or a Mann-Whitney non-parametric test for those failing the normality test as noted (TNF and IL1A versus PBS). These data were analyzed using GraphPad Prism 9 software. Following assessment, significance was noted using the following parameters: *= p<0.05, **= p<0.01, ***= p<0.001, ****= p<0.0001. Roughly, equal numbers of male and female mice were used per experiment.

### Murine RNA Isolation and Sequencing

Mouse eyes were harvested at 2 days following injection and retinas were freshly dissected in ice-cold PBS and frozen on dry ice and stored at −80° C prior to homogenization at the University of Rochester. Samples were shipped overnight to the Jackson Laboratory for RNA sequencing studies. Tissues were homogenized in MR1 buffer (Macherey-Nagel) using a Pellet Pestle Motor (Kimbal). Total RNA was isolated from tissue using the NucleoMag RNA Kit (Macherey-Nagel) and the KingFisher Flex purification system (ThermoFisher) as per the manufacturer’s protocol. An additional DNase treatment was done using the RNeasy Mini kit (Qiagen). RNA concentration and quality were assessed using the Nanodrop 8000 spectrophotometer (Thermo Scientific) and the RNA 6000 Nano Assay (Agilent Technologies). Libraries were constructed using the KAPA mRNA HyperPrep Kit (Roche Sequencing and Life Science), according to the manufacturer’s protocol. The quality and concentration of the libraries were assessed using the D5000 ScreenTape (Agilent Technologies) and Qubit dsDNA HS Assay (ThermoFisher), respectively, according to the manufacturers’ instructions. Libraries were sequenced 150 bp paired-end on an Illumina NovaSeq 6000 using the S4 Reagent Kit v1.5.

### Murine RNA-Seq Data Analysis

The resulting ~40 M read pairs per sample were processed following standard quality control practices and high-quality read pairs were aligned to the mouse genome (mm10) using STAR 2.7.9a ^36, 37^. Read pair counts per gene were summed with the featureCounts function in subread 2.0.1 ^38, 39^. Read counts were normalized using the trimmed mean of M values (TMM) method in edgeR 3.36.0 within the R environment 4.1.3 ^40, 41^. Data was scaled using the limma v3.50.3 voom function and to control for repeated measures effects, the DuplicateCorrelation function was used to block on “mouse ID” ^42^. Linear models were utilized to identify differentially expressed genes fit by the predictors of the model: Sex and type of treatment (IL1, TNF, IL1+TNF, or PBS). For the RNA-seq data involving *Sarm1* mice we also utilized the data generated for IL1, TNF, IL1+TNF treated eyes by incorporating a “batch” term into the linear model as written: Group (genotype and treatment combination) + Sex + Batch. Significant differentially expressed genes were determined by a cut-off of adjusted *p* value < 0.05 and subjected to over representation analysis of KEGG pathways using the R package clusterProfiler v4.2.2 ^43, 44^. Significant differentially expressed genes with false-discovery rate (FDR) < 0.05 and corresponding log_2_ fold changes were used as input for ingenuity pathway analysis (IPA) upstream regulator analyses (QIAGEN Inc.)^45^. KEGG pathway maps were downloaded and colored by log_2_ fold change using the R package pathview v 1.34.0 ^46^. KEGG network graphs were generated using clusterProfiler v4.2.2 ^47^. Heatmaps were generated by using the average group log (counts per million mapped reads + 1) and scaling across each gene in which the scaled value = (original value - µ) / σ via pheatmap v1.0.12. Adjustments were made for multiple testing to control the FDR at 0.05 ^48^. Data will be made available through the Gene Expression Omnibus (GEO accession pending).

## Results

### IL1A+TNF in combination, but not alone, is sufficient to drive RGC death

Previous studies have shown that intravitreal injection of TNF drives approximately 15-20% loss of RGC ^11, 12, 13^, however this death is only present 8-12 weeks following insult. Despite previous studies implicating IL1A in glaucomatous neurodegeneration, the *in vivo* role of IL1A in facilitating RGC death has not been assessed. To determine the direct neurotoxicity of IL1A on RGC viability, eyes of C57BL/6J mice were intravitreally injected with IL1A alone (0.1μg/μl). Twelve weeks after IL1A injection, there was no significant loss of RGCs compared to PBS-injected contralateral control eyes (Fig. 1A). Twelve weeks was chosen as this is a timepoint where TNF insult had previously been reported to show RGC loss ^11, 12, 13^ and this confirmed in our study (Fig. 1A). To determine if combined injection of IL1A+TNF resulted in increased RGC loss, ILA and TNF were injected together. Co-injection did not result in any increased cell death, compared to TNF injection alone 12 weeks post-injection (Fig. 1A). While TNF or IL1A injection alone did not cause significant RGC death 14 days post-injection ^12^, combined injection of IL1A+TNF did (Fig 1B-D).

**Figure 1.**
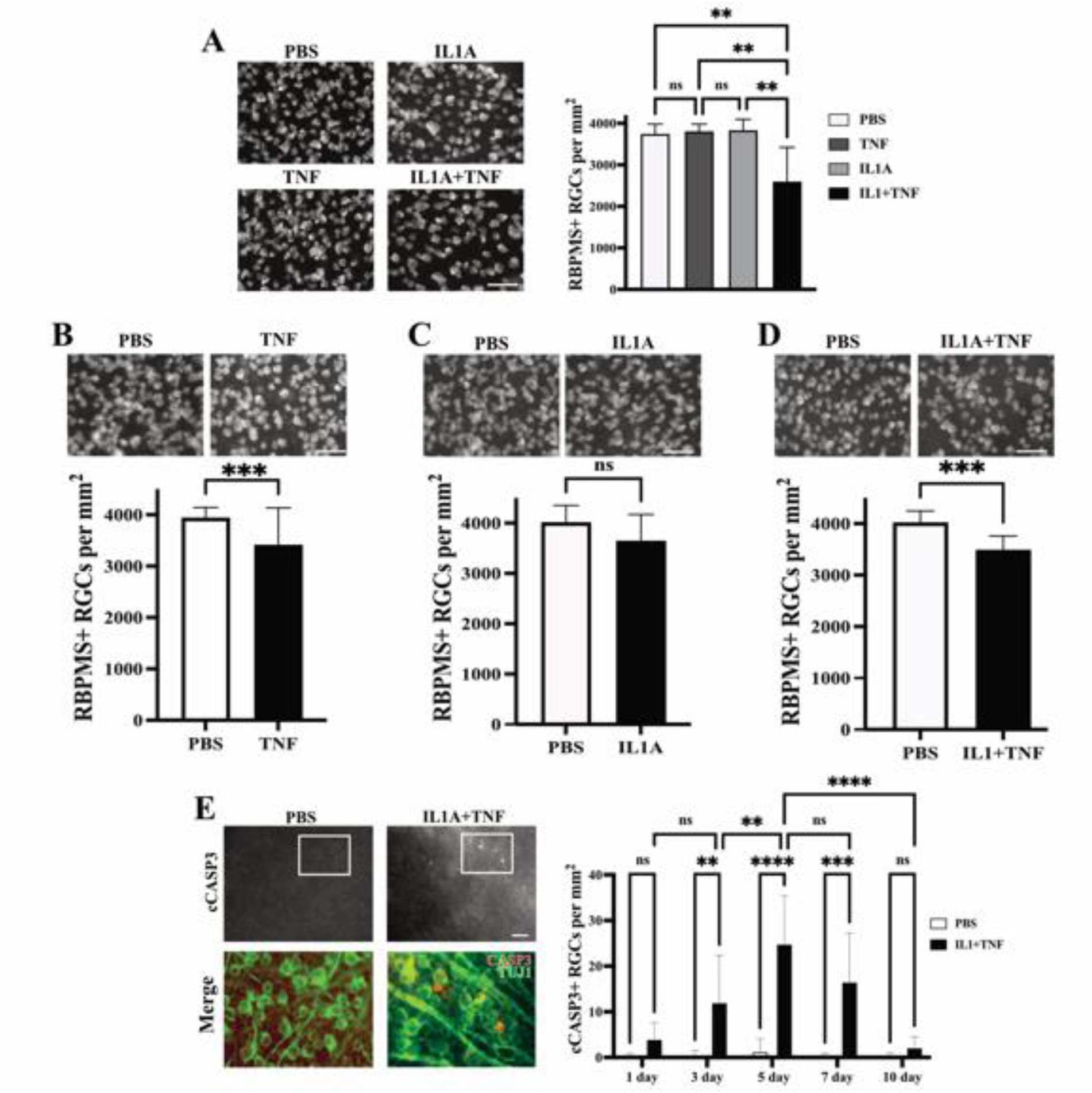
Intravitreal injection IL1A+TNF combined, but not alone, drives rapid RGC death. **A.** 14 days after intravitreal application of IL1A or TNF alone, there was no loss of RGCs as judged by RBPMS+ RGCs. However, combined application of IL1A+TNF resulted in significant (P<0.01) loss compared to PBS (control), IL1A, or TNF injection (PBS, n=26, TNF, n=7, IL1A, n=9, IL1A+TNF, n=10; **\*\***p<0.01; ns, not significant). The long-term effect of TNF, IL1A, and IL1A+TNF was investigated at 12 weeks post intravitreal injection (**B-D**) **B.** At this time point, TNF injection resulted in a significant loss of RBPMS+ RGCs compared to PBS-injected eyes (P<0.001; n=13 and 11, respectively). **C.** In contrast, IL1A injection did not result in RGC loss compared to PBS injection (n=8 for both conditions, P=0.12). **D.** However, the combined application of IL1A+TNF resulted in significant RGC loss compared to PBS (P<0.001; N=10 for both conditions). **E.** Cleaved caspase 3 (cCASP3) immunolabeling (red) was assessed in RGCs (colocalization with TUJ1+ cells (green)) at 1,3,5,7, and 10 days following IL1A+TNF or PBS application. cCASP3+TUJ1+ cell counts (per mm^2^) show a significant increase in cCASP3+TUJ1+ cells starting at around 3 days after injection and ending between 7 and 10 days (n ≥ 6 per group). Note: merged images in the bottom row indicate a zoomed-in section of the boxed area in the top row to highlight overlap. Statistical information: **A**, Kruskal Wallis Test with Dunn’s Multiple Comparisons Test, **B-D**, Brown Forsythe and Welch one way ANOVA, **E** two-way ANOVA; NS, not significant; **p<0.01, ***p<0.001, ****p<0.0001; error bars, SEM; scale bar = 50 μm.

To define the time window when RGCs die following IL1A+TNF injury, IL1A+TNF were injected into retinas, which were assessed for cleaved caspase 3 (cCASP3; a marker of dying cells). at 1,3,5,7, and 10 days. TUJ1+ cCASP3+ cells were present in the IL1A+TNF, but not PBS-injected eyes starting at 3 days and peaking between 5- and 7-days post-injection (TUJ1 labels RGCs in the ganglion cell layer; Fig. 1E). These data show that IL1A or TNF alone are not capable of driving death at early timepoints, but together can mediate rapid RGC death.

### IL1A+TNF facilitates increased neuroimmune transcriptional signatures and activation of cell-death pathways

The molecular pathways leading from TNF or IL1A+TNF insult to RGC death are unknown. It is possible these cytokines act directly on RGCs as receptors for both molecules are expressed on RGCs. It is also possible that the effect of these molecules is extrinsic to RGCs, perhaps through astrocytes, as suggested by the work of Liddelow and colleagues ^4, 5, 7^. To understand how IL1A and/or TNF drive RGC death, RNA-sequencing was performed on whole retinas isolated two days post intravitreal injection, the timepoint when RGC loss was first observed after IL1A+TNF injection. RNA-seq was performed on retinas injected with IL1A, TNF, or IL1A+TNF and contralateral control eyes injected with PBS. The results indicate that while IL1A or TNF alone induced significant transcriptional changes, the largest changes were observed in the IL1A+TNF group (Fig. 2A-D). Overrepresentation analyses revealed key pathways affected in the combined treatment group included apoptosis, necroptosis, TNF signaling, Toll-receptor signaling, and NOD-like signaling (Fig. 2E-G). Thus, the RNA-seq data revealed potential pathways important in cell signaling pathways intrinsic to RGCs and RGC extrinsic inflammatory and/or glial activation pathways. Note, when investigating the effects of TNF alone at 2, 14, and 35 days post-injection, few transcriptional changes were seen at 2 days compared to PBS controls and no significant changes at 14 or 35 days (Figure 2A, not shown).

**Figure 2.**
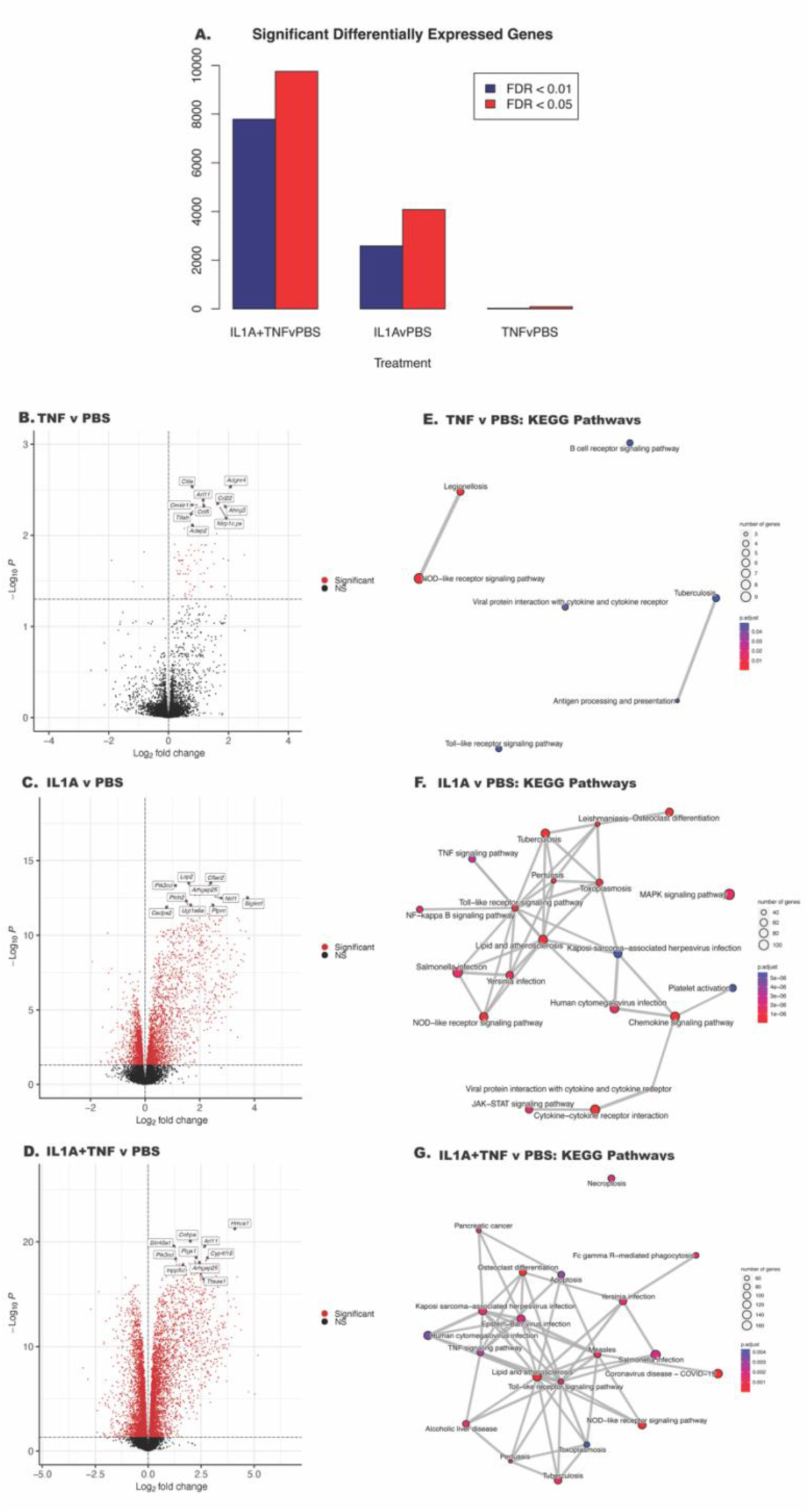
IL1A+TNF injection alters neuroinflammatory and cell death-associated transcriptional signatures. **A.** Bar chart summarizing the number of differentially expressed genes (DEGs) between the following groups from left to right: IL1A+TNF vs PBS, IL1A vs PBS, and TNF vs PBS. Red bars represent DEGs with an FDR <0.05, and dark blue bars are DEGs with an FDR < 0.01. **B-D**. Volcano plots of differential expression between **(B)** TNF vs PBS, **(C)** IL1A vs PBS, and **(D)** IL1A+TNF vs PBS. Genes with FDR < 0.05 are colored red. **E-G**. Enrichment plots of enriched KEGG gene sets between: **(E)** TNF vs PBS, **(F)** IL1A vs PBS, and **(G)** IL1A+TNF vs PBS. Lines between nodes reflect shared genes. Node size represents the number of DEGs in the gene set. The color is an illustration of the adjusted p-value for each gene set. Number of retinas per group: 31 PBS (15 female, 16 male), 8 TNF (4 female, 4 male), 9 IL1A (4 female, 5 male), and 10 IL1A+TNF (6 female, 4 male).

The transcriptional states of microglia and astrocytes have been suggested to be important for RGC loss in various disease models, particularly in association with neurotoxic cytokines ^4, 5, 7^. Genes associated with “A1” and “A2” astrocytes ^4, 49^ (Fig. 3A) were examined as well as several genes relevant to previously defined microglial disease-relevant states ^50^ (Fig. 3B). IL1A alone, but not TNF, resulted in a broad increase in astrocyte activation markers, while IL1A+TNF together was sufficient to facilitate increased expression in pan reactive, “A1”, and “A2” astrocyte markers. IL1A+TNF also led to an upregulation of gene signatures for interferon-responding microglia (IRM) and disease-associated microglia (DAM), further supporting their capability in driving glial activation 2 days post-injection.

**Figure 3.**
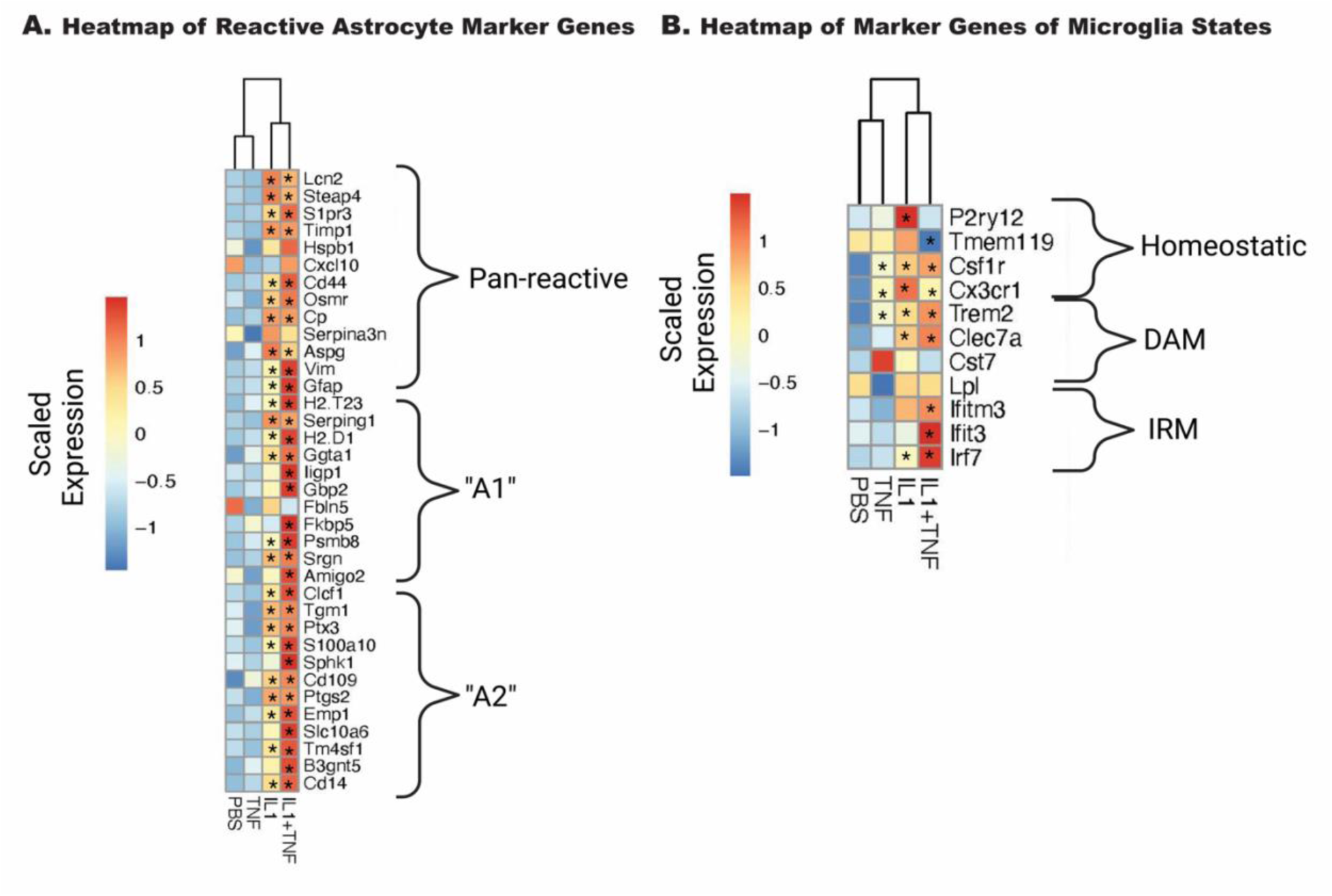
IL1A+TNF induces transcripts associated with glial activation. Heatmaps of curated genes associated with astrocytes (**A**) and microglia (**B**). Average expression for the entire group is shown. Scaled and centered expression of the library-size and log normalized average expression value for each gene in each treatment group. * denotes FDR <0.05 for the selected gene in the treatment group when compared to the PBS group; numbers retinas per group: 31 PBS (15 female, 16 male), 8 TNF (4 female, 4 male), 9 IL1A (4 female, 5 male), and 10 IL1A+TNF (6 female, 4 male).

### IL1A+TNF induced intrinsic RGC death pathways

We were particularly interested in understanding the molecular pathways within RGCs following cytokine insults. Overrepresentation analysis showed the apoptosis pathway was significantly overrepresented in differentially expressed genes comparing IL1A+TNF with PBS retinas at 2 days after injection (Fig. 4A). Previous work has shown that transcription factors can be critical mediators of RGC death after glaucoma-relevant injury ^31, 32, 35, 51, 52^. To identify putative master regulators of transcriptional changes identified in the IL1A+TNF group, we utilized the Upstream Regulator Analysis tool in Ingenuity Pathway Analysis (IPA). Upstream regulator analysis suggested the activation of several transcriptional regulators, including JUN (Fig. 4B).

**Figure 4.**
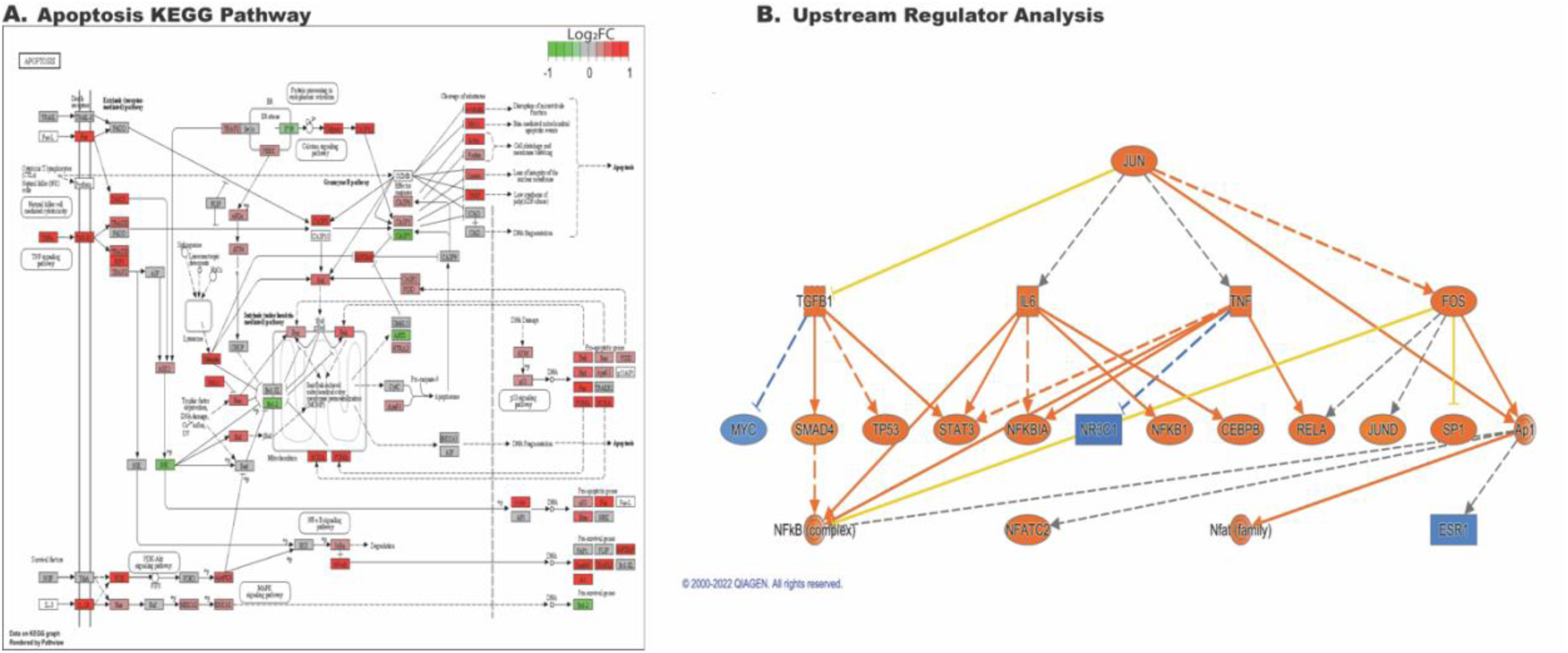
IL1A+TNF co-treatment induces cell-death and activation of JUN transcriptional signatures. **A.** KEGG graph of the significantly enriched Apoptosis pathway within the DEGs comparing IL1A+TNF vs PBS and is colored by the logFoldChange value for the expression of highlighted genes within the pathway comparing IL1A+ vs PBS. **B.** IPA upstream regulator analysis generated a mechanistic network for JUN using a strict cut-off for input DEGs identified between IL1A+TNF vs PBS with FDR < 0.01 and LogFC > 0.5. Orange and blue indicate predicted activation and inhibition, respectively. Solid and dashed lines, respectively, indicate direct and indirect interactions. The yellow line suggests inconsistent with the state of downstream molecules. Gray indicates no effect predicted. Number of retinas used per group: 31, PBS (15 female, 16 male), 8 TNF (4 female, 4 male), 9 IL1A (4 female, 5 male), and 10 IL1A+TNF (6 female, 4 male).

JUN is a transcription factor involved in injury response in RGCs. Furthermore, JUN is important for RGC death after several glaucoma-relevant injuries, including OHT and ONC ^31, 32, 35, 51^. Given the enrichment of JUN in the upstream regulator analysis and its known importance in RGC death, we assessed whether JUN is activated (phosphorylated; pJUN) downstream of intravitreal IL1A+TNF insult in RGCs. Three days after IL1A+TNF injection, before peak RGC degeneration, pJUN was present in RBPMS+ cells, and absent in PBS injected eyes (Fig. 5A). Thus, JUN is activated in RGCs after IL1A+TNF insult. To determine if JUN is required for RGC death, mice deficient in *Jun* in RGCs and macroglia were generated using floxed alleles of *Jun* ^29^ (*Jun^fl^*) recombined with *Six3*-cre ^30^. *Six3*-cre^+^*Jun^fl/fl^* and control mice (e.g., *Six3*-cre^+^*Jun^fl/fl^*or *Six3*-cre^−^ *Jun^fl/+^*) were intravitreally injected with either IL1A+TNF or PBS. Contrary to the protective effects of the loss of *Jun* seen in other glaucoma-relevant injuries, ^31, 32, 33, 51^, *Six3*-cre^+^*Jun^fl/fl^*animals showed no protection to RGCs from IL1A+TNF insult (Fig. 5B). These findings suggest that while JUN is activated after IL1A+TNF injury, RGC death is independent of *Jun* activation.

**Figure 5.**
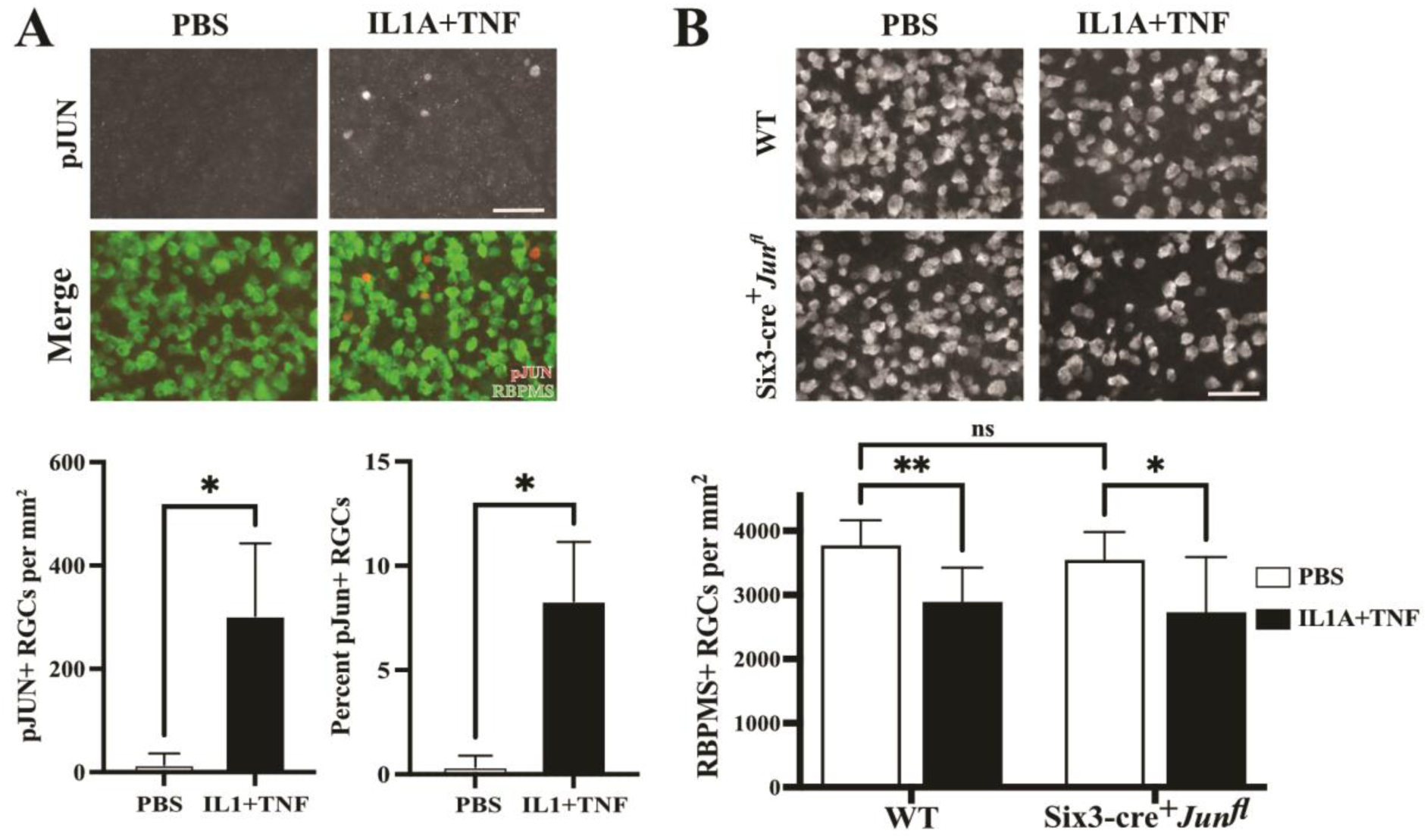
While JUN is activated following IL1A+TNF, *Jun* expression in RGCs and macroglia is not required for IL1A+TNF mediated degeneration. **A.** 3 days post IL1A+TNF injection, phosphorylated c-Jun (pJUN) is expressed in significantly more RBPMS+ RGCs compared than in control eyes injected with PBS (given as pJUN+ RGCs per mm^2^; PBS median, 5.0; IL1A+TNF median; 250.8; *p=0.028, n=4 per condition, Mann Whitney test). This equates to approximately 8% of RGCs after IL1A+TNF injection and 0.2% of RGCs after PBS injection (*p=0.028, n=4 per condition, Mann Whitney test). **B.** Following IL1A+TNF injection, animals with *Jun* deleted in RGCs and macroglia had a loss of RGCs compared to PBS control eyes (RBPMS+ RGCs per mm^2^± SEM: Control PBS, 3782 ± 120; Control IL1A+TNF, 2754, ± 159, n=11; *Jun−/−* PBS, 3478 ± 141; *Jun−/−* IL1A+TNF, 2739± 271, n=10; Control PBS vs Control IL1A+TNF **p<0.01; *Jun−/−* PBS vs *Jun−/−* IL1A+TNF * p<0.05; Control PBS vs *Jun−/−* PBS NS, p=0.815; Two-way ANOVA with Tukeys Multiple Comparisons test).

In the CNS, previous studies identified SARM1 (sterile alpha and TIR motif containing protein 1) as a key regulator in axonal degeneration ^53, 54, 55, 56^. While deficiency of *Sarm1* led to reduced axonal degeneration after glaucoma-relevant injury, it failed to protect RGCs somas from apoptosis and death ^53^. However, SARM1 has recently been linked to neuronal survival after TNF insult, with loss of *Sarm1* conferring near complete protection to RGC somas following TNF injection ^12^. Therefore, to assess the potential role of SARM1 following the combined application of IL1A+TNF, eyes from *Sarm1^−/−^* and WT mice were injected IL1A+TNF and RGC density was quantified 14 days after injury. In line with the RGC protection seen in *Sarm1^−/−^* animals injected with TNF ^12^, we found that *Sarm1* deficiency prevented the early IL1A+TNF-induced death (Fig. 6).

**Figure 6.**
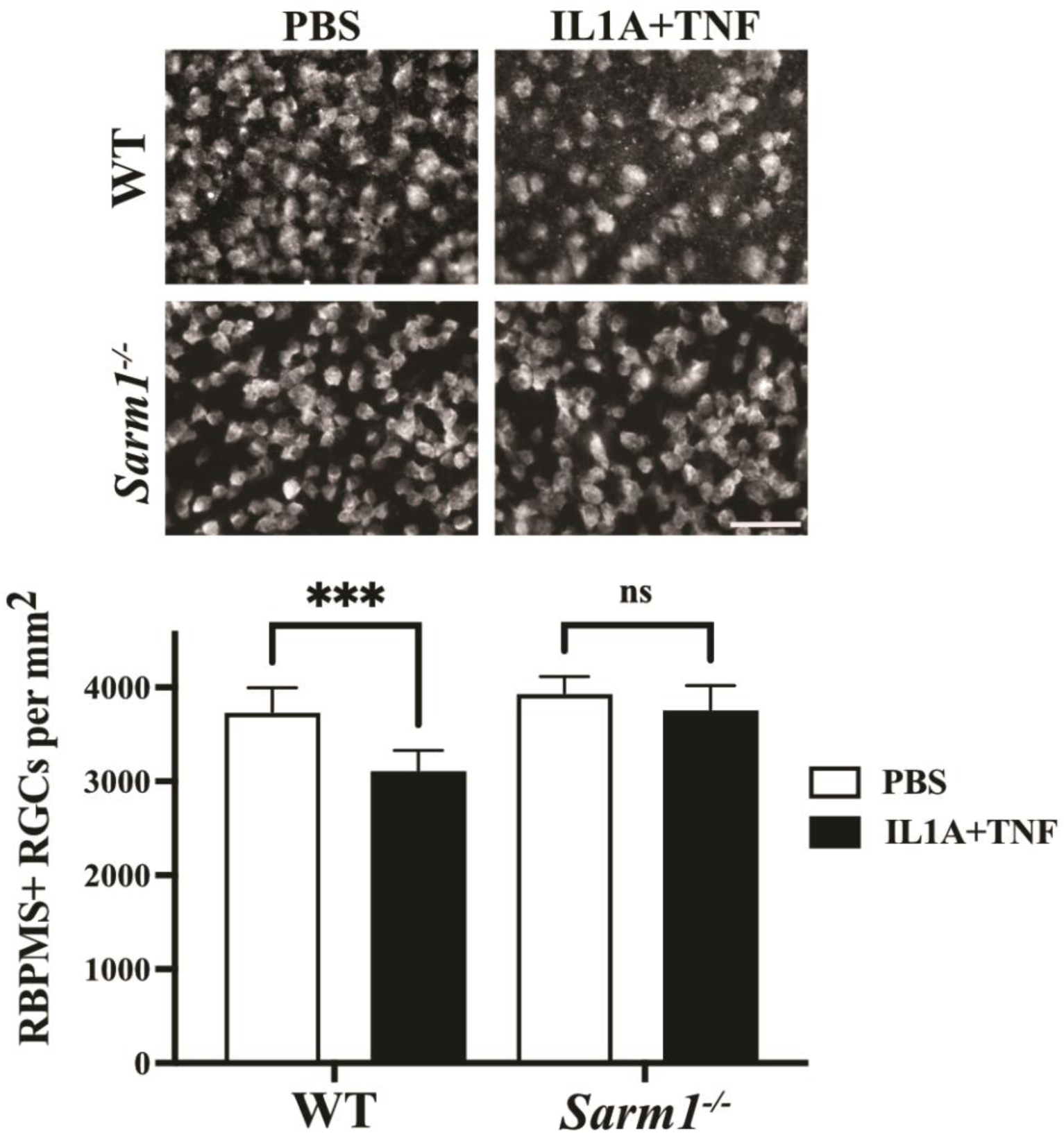
Loss of *Sarm1* expression was sufficient to prevent IL1A+TNF induced death. *Sarm1* deficient animals no RGCs loss 14 days after IL1A+TNF injection compared to PBS injected eyes (RBPMS+ RGCs per mm^2^ ± SEM; n=8 per group; p>0.05). In contrast, wildtype animals injected had significant RGC loss 14 days after IL1A+TNF, but not in PBS controls (n=9 per group; p<0.001, ***. Brown-Forsythe and Welch ANOVA were used for all comparisons; scale bar = 50 μm.

### Loss of *Sarm1* does not prevent IL1A+TNF-induced glial activation transcriptional signatures

To gain insight into the molecular mechanism of SARM1 in RGC death after IL1A+TNF insult RNA-Seq was performed on WT IL1A+TNF, WT PBS, *Sarm1^−/−^*IL1A+TNF, and *Sarm1^−/−^* PBS retinas 2 days after injections. We included samples from Figure 2 in this analysis to allow us to compare to TNF and IL1A sole treatments by including a batch term in the model. *Sarm1* deficiency without insult only resulted in 8 genes being differentially expressed in uninjured eyes: *Sarm1*, *Slc46a1*, *Dynlt1b*, *Fam57a*, *Vsp53*, *Tlcd2*, *ENSMUSG00000079733*, and the pseudogene *Tmem181b.ps* (Figure 7A-B). There was differential expression of genes associated with synaptic changes, calcium signaling, MAPK signaling, and glial responses between *Sarm1* deficient and control retinas (Figure 7C-F) after IL1A+TNF insult. Yet, *Sarm1* deficient animals had similar levels of astrocyte activation (Figure 8A) and microglial response (Figure 8B) 2 days after IL1A+TNF insult. Therefore, while *Sarm1* deficiency can prevent IL1A+TNF induced RGC loss, it is not necessary for IL1A+TNF mediated glial activation.

**Figure 7.**
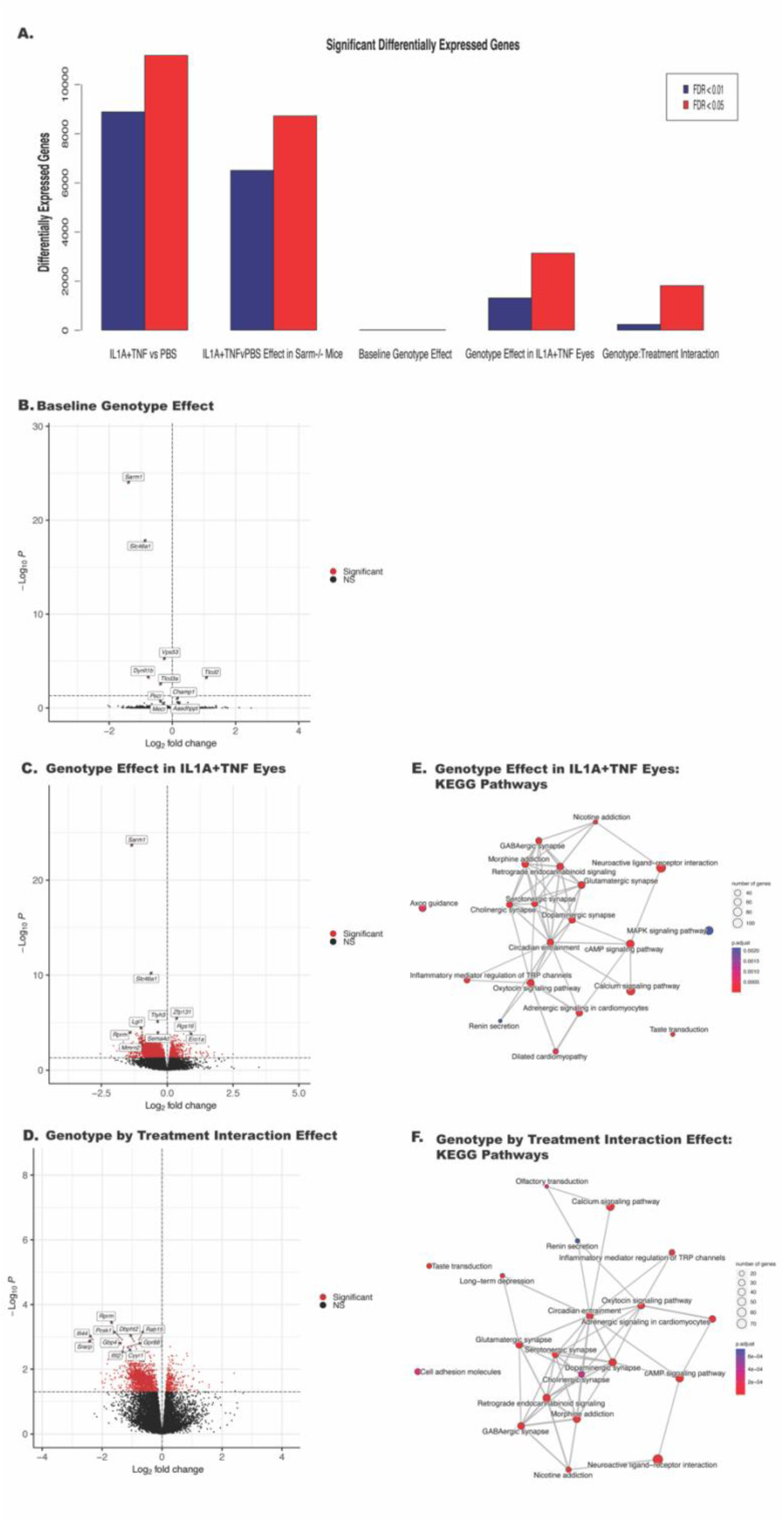
*Sarm1* deficiency significantly alters synapse-associated transcripts after IL1A+TNF insult. **A.** Bar chart summarizing the number of differentially expressed genes (DEGs) between the following groups from left to right: IL1A+TNF vs PBS in WT eyes, IL1A+TNF vs PBS in *Sarm1^−/−^* eyes, *Sarm1^−/−^*retinas vs WT retinas, IL1A+TNF treated *Sarm1^−/−^* vs IL1A+TNF WT eyes, and the Genotype by Treatment interaction effect. Red bars represent DEGs with an FDR <0.05 and dark blue bars are DEGs with an FDR < 0.01. **B-D**. Volcano plots of differential expression between (**B**) *Sarm1^−/−^* retinas vs WT retinas, (**C**) IL1A+TNF treated *Sarm1^−/−^* vs IL1A+TNF treated WT eyes, and (**D**) Genotype by Treatment, genes with FDR < 0.05 are colored red. **E-F**. Enrichment plots of enriched KEGG gene sets between: (**E**) IL1A+TNF treated *Sarm1^−/−^* vs IL1A+TNF treated WT eyes, and (**F**) Genotype by Treatment. Lines between nodes reflect shared genes and node size represents the number of DEGs in the gene set. The color is an illustration of the adjusted p-value for each gene set. Retinas used by treatment: PBS, 36 WT (17 female, 19 male) and *6 Sarm1^−/−^* (3 female, 3male); IL1A+TNF, 16 WT (9 female, 7 male) and 6 *Sarm1^−/−^* eyes (3 female, 3 male).

**Figure 8.**
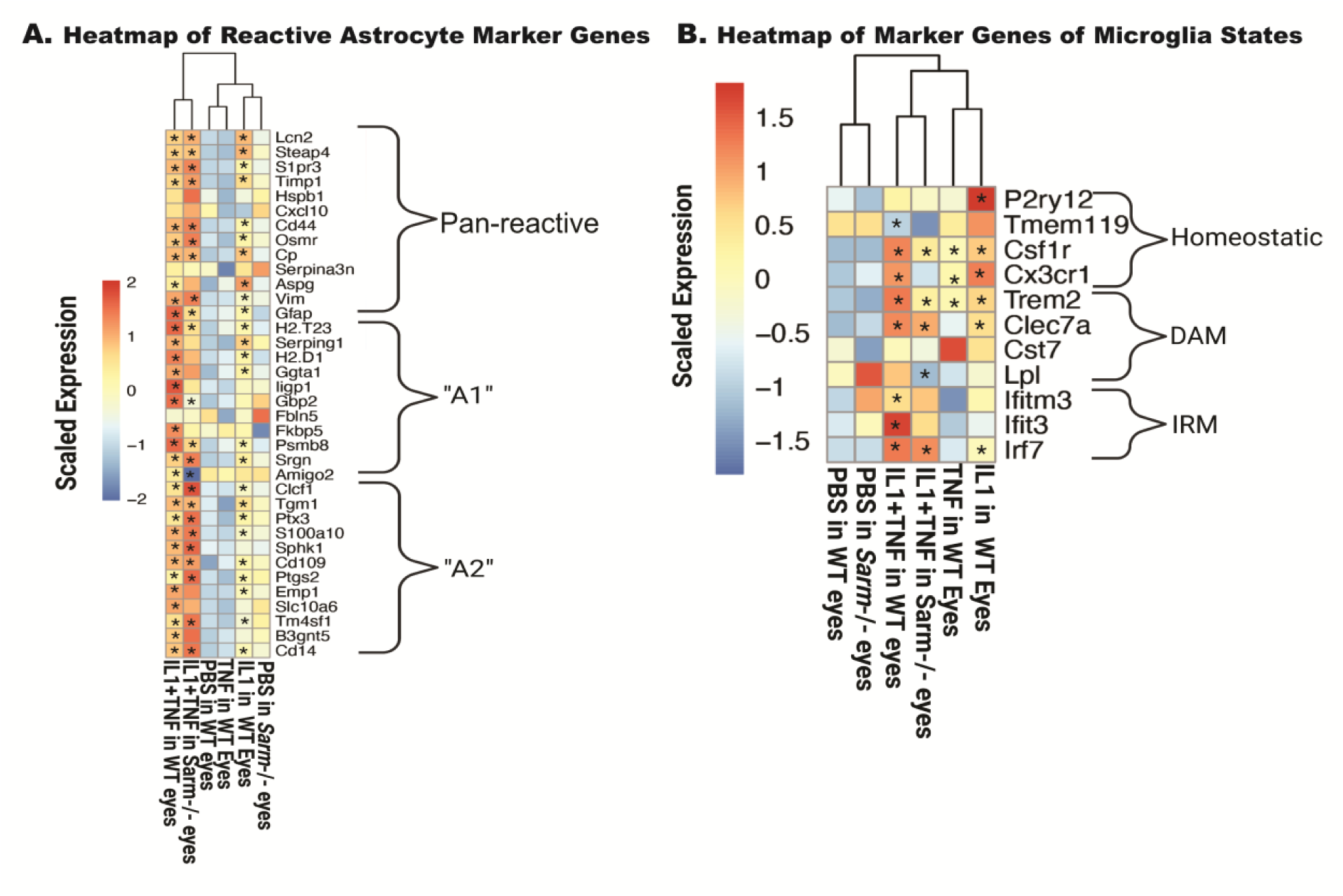
*Sarm1* deficiency does not reduce glial activation driven by IL1A+TNF. Heatmaps of curated genes associated with astrocytes (**A**) and microglia (**B**). Average expression for the entire group is shown. Scaled and centered expression of the library-size and log normalized average expression value for each gene in each treatment group is shown. * denotes FDR <0.05 for the selected gene in the treatment group when compared to the PBS in WT eyes or PBS in *Sarm1*^−/−^ eyes. Retinas used by treatment: PBS, 36 WT (17 female, 19 male) and *6 Sarm1^−/−^* (3 female, 3male); IL1A+TNF, 16 WT (9 female, 7 male) and 6 *Sarm1^−/−^* eyes (3 female, 3 male).

## Discussion

Neuroinflammation has been extensively linked to RGC cell death and glaucomatous neurodegeneration ^8, 57, 58, 59, 60^. Recent studies have shown that combined deficiency of the neuroinflammatory-related molecules *Tnf*, *C1q*, and *Il1a* prevented RGC death after ONC ^4, 5^ and lessened RGC death after an OHT injury ^6^. While TNF and C1Q have been studied extensively in glaucoma-relevant pathogenesis, IL1A has not been assessed for its neurotoxic capabilities to RGCs *in vivo*.

Intravitreal injection of TNF results in approximately 15-20% loss of RGCs, however, the loss of RGCs appears only around 8-12 weeks following injection ^11, 12, 13^. IL1A has been assessed for its ability to drive neuroinflammation in the brain ^61, 62^, but to our knowledge, its effect on the retina in vivo has never been tested. To determine the direct neurotoxicity of IL1A and the combinatorial effects of IL1A+TNF, the eyes of B6 mice were intravitreally injected with IL1A alone, TNF alone, or IL1A+TNF. Consistent with previous studies ^11, 63^, TNF injection failed to kill RGCs by 14 days following insult but did result in RGC loss 12 weeks post-injection. In addition, IL1A alone did not result in any loss of RGCs at any time point assessed out to 12 weeks (the latest time point examined). However, in eyes receiving a combined injection of IL1A+ TNF, there was a significant decrease in RGC density at 14 days (~20%) compared to control eyes, with RGCs dying mainly between 3 and 7 days. Interestingly, at 12 weeks, the amount of death remained ~20%, suggesting IL1A acted as a sensitizer for TNF-induced RGC death, accelerating RGC death, but did not drive RGC death on its own.

The molecular pathways leading from TNF or IL1A+TNF insult to RGC death is unknown. These cytokines could act directly on RGCs as their receptors are expressed on RGCs. As Liddelow and colleagues ^4, 5, 7^ have suggested, it is possible that TNF and IL1A act on astrocytes, which in turn release a neurotoxic substance. To gain insight into this question, RNA-seq on whole retinas was performed 2 days following either TNF, IL1A, or IL1A+TNF insult. This time point was chosen to identify primary changes after cytokine insult. Only around 50 genes had significant changes in their expression after TNF insult alone. Also, surprisingly, TNF injection did not result in any significant gene expression changes at either 14 or 35 days. Thus, these experiments did not shed any insight on why TNF induces a delayed RGC death. In contrast to TNF, intravitreal injection of either IL1A alone or IL1A+TNF led to significant transcriptional changes relative to PBS controls (2000+ and 8000+ genes, respectively). Pathway analysis revealed several interesting pathways to be upregulated, suggesting possible mechanisms for IL1A+TNF-induced RGC death. These included both apoptosis and necroptosis, as well as TNF, NOD-like, and Toll-like receptor signaling pathways. These findings demonstrated the early capability of these cytokines to drive both RGC death and immune-related signaling changes by 2 days.

We were particularly interested in whether glial activation occurred prior to or coincidently with RGC death after IL1A and IL1A+TNF injections. In OHT models of glaucoma, glial activation occurs before RGC loss ^8, 64^. After mechanical axon injury to RGCs, our group showed using RNA-seq data that proinflammatory signaling and glial activation in the retina might be secondary to activation of pro-death pathways in RGCs after acute axonal injury ^52^. However, Liddelow and colleagues have shown that cytokine signaling kills RGCs, assumed to be acting through astrocytes ^4, 7^. Our RNA-seq data here showed that marker genes for astrocyte and microglial responses were upregulated. For microglia, expression changes showed clear upregulation of both IRM and DAM microglial responses. There were also significant changes in genes that are important in astrocyte activation. Upregulation of genes thought important in astrocytic states associated with pathology were present, including genes marking A1, A2, and pan reactive astrocytes. Interestingly, unlike previous *in vitro* findings^4^, IL1A+TNF insult appeared to mediate not just A1, but also A2 signatures, suggesting that *in vivo* these cytokines do not act solely in an A1-mediated neurotoxic manner. However, because we performed bulk RNA-seq, we do not know if these marker genes were upregulated in distinct cell (sub)populations. Together, these data show that both microglia and astrocytes are activated at the onset of RGC injury and are consistent with an important role for glia in IL1A+TNF-induced RGC death.

Along with the role of neuroinflammatory signaling in IL1A+TNF-induced RGC death, we were also interested in understanding cell signaling pathways within RGCs that lead to RGC death after IL1A+TNF insult. The RNA-seq analysis showed that cell death pathways, including apoptosis and necroptosis, were overrepresented. It was particularly interesting that upstream regulator analysis of the differentially regulated transcripts identified the transcription factor JUN. JUN is important in RGC death after axonal injury and in ocular hypertensive eyes ^31, 32, 35, 51, 52^.

To determine the downstream signaling events following IL1A+TNF insult before RGC death, we assessed animals injected with IL1A+TNF at 3 days for phosphorylated-c-jun (pJUN), a known transcription factor involved in RGC death following a variety of insults. At this time point, JUN is significantly elevated in IL1A+TNF injected retinas compared to PBS controls, with approximately 8-10% of RGCs expressing pJUN (Figure 6A). These findings indicated that JUN signaling is occurring downstream of IL1A+TNF insult. However, to assess whether *Jun* is required for RGC death after this insult, we recombined *Jun^fl^* alleles using *Six3*-cre to target RGCs and macroglia. Interestingly, loss of Jun in Six3-cre^+^ *Jun^fl^* animals failed to prevent RGC death after IL1A+TNF injection (Figure 2.6B). These findings contrast with other glaucoma relevant insults showing protection to RGCs following loss of *Ju*n ^31, 32^, suggesting a *Jun* independent mechanism of RGC death following extrinsic application of IL1A+TNF. The RNA-seq pathway analysis also identified necroptosis as a potential cell death pathway acutely activated following IL1A+TNF. SARM1 is involved in necroptosis ^12, 65^. Furthermore, SARM1 has recently been linked to neuronal survival after TNF insult, and, importantly, SARM1 is known to be involved in axonal and degeneration after axonal insult ^53, 54, 55, 66^, a key component of glaucomatous neurodegeneration. Ko and colleagues recently also found that *Sarm1^−/−^* mice showed no RGC somal or axonal loss following TNF injection ^12^, suggesting SARM1 may play a key role in RGC somal death after a neurotoxic insult. Not surprisingly, *Sarm1^−/−^*mice showed no loss of RGCs following IL1A+TNF (Figure 7), supporting the role of SARM1 in the soma after neurotoxic cytokine insult. Interestingly, SARM1, while typically described as a modulator of NAD+ metabolism and axon integrity, is also known as Myd88-5, which is a bottleneck molecule downstream in IL1A signaling ^67^. Our findings would also support SARM1’s role in initiating downstream mechanisms following exposure to IL1A+TNF and potentially acting upstream of JUN.

RNA-seq analysis was performed on *Sarm1* deficient mice 2 days following IL1A+TNF insult. This analysis was done to gain insight into where in the cell signaling process SARM1 was acting. When assessing the differences in gene expression IL1A+TNF treated eyes across genotypes, most of the differentially expressed genes fell under synaptic signaling pathways, with the largest genotype effect seen in *Sarm1* expression itself (Figure 9C-F). These data underscore that RGC death in control eyes treated with IL1A+TNF starts at 3 days post-injury, and changes to synaptic structure likely occur before overt neuronal loss and is seemingly modulated by SARM1 deficiency. In line with this, and in contrast to WT-treated eyes, SARM1-deficient eyes displayed an increased expression of *Nrf1*, previously implicated in the survival of developing RGCs and regulates mitochondrial function ^68^.

When comparing astrocyte activation markers, there appears to be a subtle but significant reduction in A1, but not A2 transcripts, in *Sarm1^−/−^*treated eyes (Figure 8A). When assessing changes in microglial-associated transcripts, we saw similar levels of IRM and DAM transcript upregulation in both WT and *Sarm1^−/−^* eyes after IL1A+TNF (Figure 8B). These results indicate that there are unique transcriptional differences (~2000 genes) following IL1A+TNF when cell death is prevented via loss of *Sarm1*, relative to WT eyes, but the majority of IL1A+TNF associated changes still occur (>8000 genes).

Overall, these findings demonstrated the direct neurotoxic role of IL1A+TNF to retinal ganglion cells *in vivo*. We determined that while RGC loss after IL1A+TNF is present by 14 days, this degeneration does not occur via the same pathways that govern RGC death after glaucoma-relevant insult. Unlike axonal or ocular hypertensive insults that require JUN for RGC somal loss, loss of Jun did not protect RGC somas from IL1A+TNF injury. Instead, loss of Sarm1 was protective, suggesting that IL1A+TNF injury does not model glaucoma-relevant injury. These data were in line with the findings of Ko et al., showing a requirement for *Sarm1* in TNF-induced RGC loss. Therefore, IL1A+TNF appears to induce a distinct form of RGC death and may serve as a model for neuroimmune mediated RGC insult, but not glaucomatous injury.

## Acknowledgments

This work was supported by the BrightFocus Foundation (RTL), National Eye Institute EY027701 (RTL, GRH), EY035093 (RTL, GRH), EY018606 (RTL), an unrestricted grant from the Research to Prevent Blindness to the Department of Ophthalmology at the University of Rochester. KMA was also supported by the Center for Visual Science Training Grant T32-EY007125 (KMA) and the Neuroscience Graduate Program at the University of Rochester. GRH is also extremely grateful for the support of the Diana Davis Spencer Foundation. We gratefully acknowledge the contribution of the Genome Technologies Scientific Service at The Jackson Laboratory for expert assistance with the work described in this publication.

## Author Contributions

KMA, GRH, and RTL designed the experiments. KMA performed all injections and immunohistochemistry experiments. MM processed and analyzed all RNA sequencing studies. All authors participated in analysis of experiments. The manuscript was written and edited by all authors.

